# A microfluidic finger-actuated blood lysate preparation device enabled by rapid acoustofluidic mixing

**DOI:** 10.1101/2022.10.16.512425

**Authors:** Md Ehtashamul Haque, Alvaro J. Conde, William N. MacPherson, Stephen R Knight, Richard M. Carter, Maïwenn Kersaudy-Kerhoas

## Abstract

For many blood-based diagnostic tests, including prophylactic drug analysis and malaria assays, red blood cells must be lysed effectively prior to their use in an analytical workflow. We report on a finger-actuated blood lysate preparation device, which utilises a previously reported acoustofluidic micromixer module. The integrated device includes a range of innovations from a sample interface, to the integration of blisters on a laser engraved surface and a large volume (130 μL) one-stroke manual pump which could be useful in other low-cost microfluidic-based point-of-care devices. The adaptability of the acoustic mixer is demonstrated on highly viscous fluids, including whole blood, with up to 65% percent volume fraction of red blood cells. Used in conjunction with a lysis buffer, the micromixer unit is also shown to lyse a finger-prick (20 μL) blood sample in 30 seconds and benchmarked across ten donor samples. Finally, we demonstrate the ease of use of the fully integrated device. Cheap, modular, but reliable, finger-actuated microfluidic functions could open up opportunities for the development of diagnostics with minimal resources.

## Introduction

Cell lysis refers to the physical, enzymatic, or osmotic mechanisms that break down the membranes of cells. Erythrocyte, or Red Blood Cell (RBC), lysis is required for downstream analysis in a variety of biological tests, including but not limited to accessing intracellular components ^1^, hydrocephalus studies ^2^, and for the isolation of human mesenchymal stromal cells ^3^. Lysis has also to be performed prior to the flow cytometry analysis of peripheral blood or certain lymphoid tissue suspensions^4^. Hybridoma protocols also necessitate the lysis of red blood cells from mouse splenocyte suspensions before fusion^5^. Many drugs, such as tacrolimus, have a high binding capacity to RBCs. Therefore therapeutic drug monitoring assays require thorough RBC lysis, prior to analysis. Other assays that require the release of intra-erythrocytic analytes include malaria assay. A large number of microfluidic devices have functional elements for blood handling, and in particular several microfluidic solutions have been proposed for RBC lysis. A paper-based microfluidic assay in conjunction with chemical hemolysis treatment has been used to quantify free-hemoglobin (Hgb) ^6^. Inertial microfluidic structures were used in work reported by Zhu et al. ^7^ and Ramachandraiah et al. ^8^ to lyse RBCs and isolate WBCs selectively. Various acoustofluidic method to lyse RBCs have been reported. In one instance, surface acoustic wave (SAW) were used to induce thermal lysis^9^. Kaba et al. have also explored a cavitation-microstreaming-based cell lysis and DNA extraction approach; however they employed Korean Cell Line Bank cells in their investigation, not blood samples^10^. More recently, Pourabed et al. proposed an acoustofluidic device capable of lysing and homogenising diluted blood samples in just 350 ms.

Notably, most of these previous microfluidic based cell lysis devices, with the exception of Baillargeon and Pourabed, need external connectors, tubing, and pumps, which makes their use laborious for simple procedures. Baillargeon paper-based solution offers an elegant simple solution but has two drawbacks: (i) it can handle only a limited volume of sample, and (ii) it has limited compatibility for downstream analytical procedures. In the case of Pourabed, despite the advantageous performance of the device, the requirement for the acoustic resonator to be manufactured into a double side polished (100) oriented silicon wafer sandwiched between two Polymethylsiloxane (PDMS) layers, makes this device difficult to reproduce and difficult to scale-up at low-cost.

In this communication, we introduce a finger-actuated blood lysate preparation device, which utilises a previously reported acoustofluidic micromixer module (Figure 1.a)^11^. Before describing the completely integrated device, which include a range of innovations from sample interface, to integration of blister on a laser engraved chip and a large volume one-stroke manual pump, we investigate the mixing performance of the acoustofluidic device onviscous fluids and human blood. Then we investigate a chemical-assisted blood lysis method and finally we detail the concept of the finger-actuated blood lysate preparation device and investigate the lysis across ten individual donor samples.

**Figure 1:**
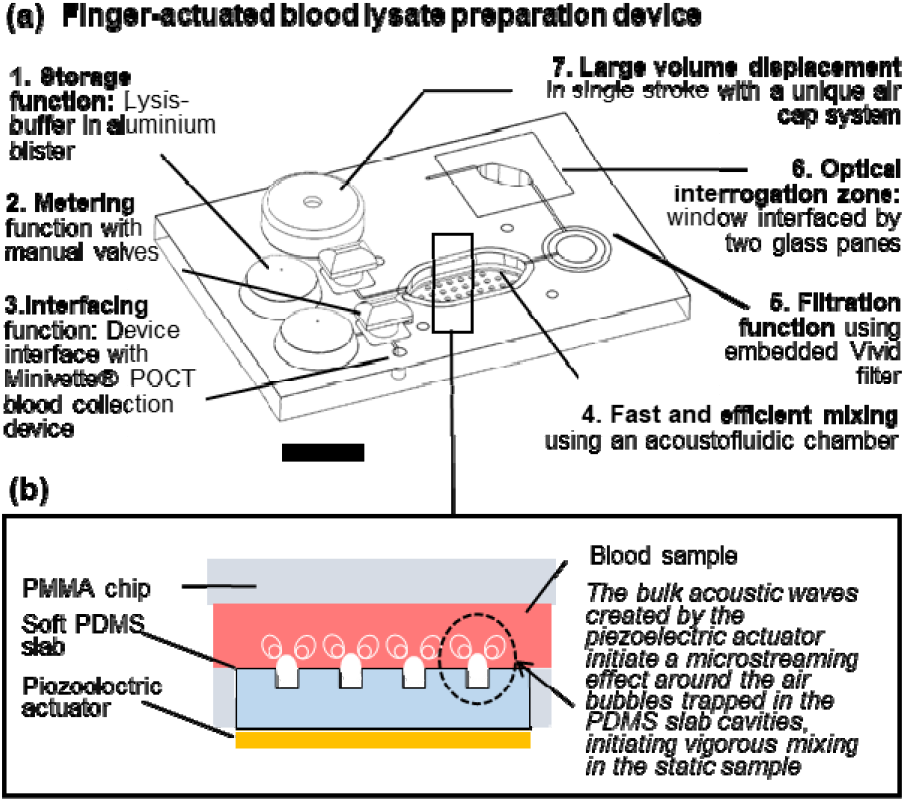
A finger-actuated filtered blood lysate preparation device (a) Isometric CAD view of the concept, illustrating seven functional elements (b) Illustration of the acoustic mixing principles developed in reference^11^

**Figure 2:**
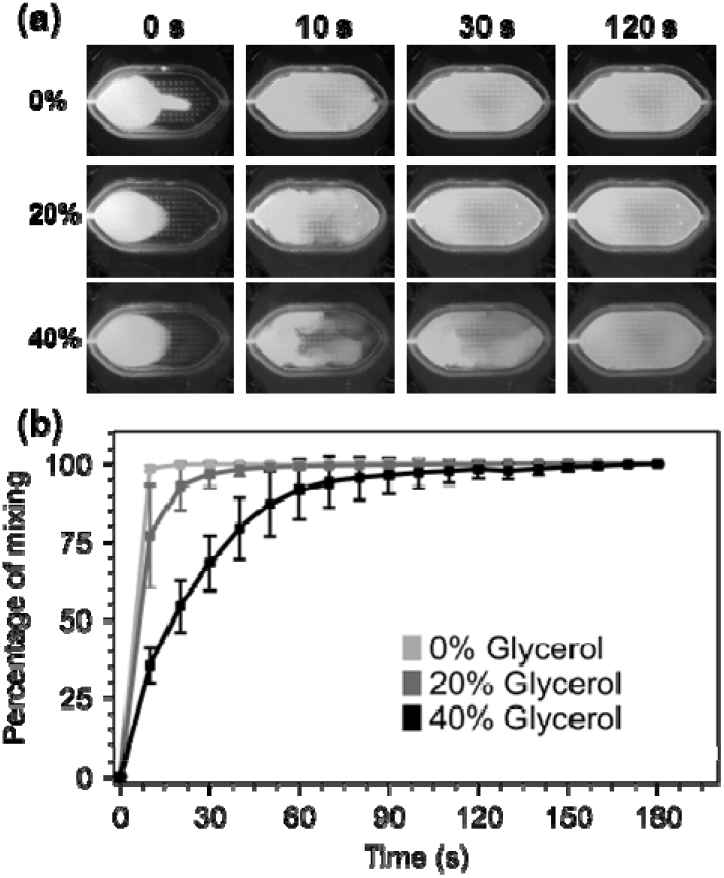
Performance of the acoustofluidic mixer with viscous fluids. a) Video screenshots of 0%, 20% and 40% glycerol solutions (Glycerol-DI water mixture was used in one half of the chamber, and Glycerol-fluorescein-tagged DI water mixture was used in the other half.) at 0 s, 10 s, 30 s and 120 s time points b) Analysis of mixing performance. Percentage of mixing was assessed by image analysis as described in materials and methods. All data acquired in triplicate.

### The acoustofluidic mixer can mix highly viscous fluids

In the absence of turbulence, or convection fluids depends exclusively on molecular diffusion to mix. Higher viscosities lower the Reynolds number and make viscous fluids especially challenging to mix in any format, including a microfluidic format. ^12–14^. Mixing performance has a direct impact on the efficiency and sensitivity of assays ^15^. Active micromixers, require external energy sources, are preferred over passive microxers due to their capacity to mix rapidly in confined volumes of liquids in the absence of flow, and thus have a smaller footprint. To date, many different types of active micromixers have been developed, utilizing a variety of external sources such as pressure ^16,17^, acoustics ^18,19^, magnetic fields ^20,21^, electric fields ^22,23^, thermal fields ^24,25^, etc. Reviews of active micromixer can be found in15,26,27.

Amongst all of these techniques, the acoustic vibration of bubbles, and resulting microstreaming effect, has become a widespread way to mix fluids due to their efficacy (fast mixing down to milliseconds). Such microstreaming acoustic mixing is often compatible with simple geometries and often utilises cost-effective actuators in the form of simple piezoelectric disks. However, most techniques reported to date involve complex prototyping methods^10^. Our team has previously demonstrated a low-cost rapid prototyping technique for fabricating a simple but agile acoustofluidic micromixer module^11^. This acoustofluidic micromixer is made up of two parts: a hard polymer cassette made of laser-cut cell-cast polymethyl methacrylate (PMMA) sheets (Clarex, various thicknesses) that house the mixing chamber and a soft polymer made of polydimethylsiloxane (PDMS), which focuses the acoustic excitation provided to the chamber by a piezoelectric actuator (Figure 1.b) ^11^. Using this device, we have reported cfDNA extraction on 100 μL plasma samples ten times superior to a standard benchmark. Here, we sought to investigate the effect of viscosity on the acoustic micromixer performance. To this effect, two types of solutions were prepared. The first solution contained either 0%, 20%, and 40% glycerol in deionised (DI) water, while the second solution contained either 0%, 20%, and 40% glycerol in DI water spiked with 5% fluorescein (v/v). 65 µL of each solution was pipetted into the mixing chamber for a specific solution experiment. After loading, a piezoelectric diaphragm (7BB-15-6L0, Murata Electronics) was driven by a function generator (TG215, Thurlby Thandar Instruments) at 4 ± 0.5 kHz and 20 Vpp.

The mixing experiment videos were recorded with digital microscopes equipped with blue LEDs for excitation and a 510 nm emission filter (AM4115T-GFBW, Dino-Lite). Sample videos are available in SI Video 1. The recorded images were processed with the ImageJ software (version 1.53q) (See SI Section 1 for full details). Mixing performance of various glycerol mixtures at various time instances can be observed from video screenshots in Figure 1.a.

As expected, the mixing performance decreases as the viscosity of the samples increases from 1 cP (viscosity of water at 20°C) to 2 cP (20% glycerol) and 4.8 cP (40%). Moving liquids of higher viscosity from one region to another requires more energy, and because the mixer operates at a fixed energy, higher viscosity results in slower mixing. Despite the micromixer’s slower mixing for highly viscous solutions full mixing of water-based solutions (0%) and 20% glycerol solutions was achieved in under 60 s while the 40% glycerol solution was mixed in under 2 min (Figure 1.b), comparable to results that have been reported in another BAW based micromixing work ^14^. However, it is inferior to other acoustic mixers reporting millisecond-mixing rates. This can be explained by the fact that the volumes mixed here are much larger than other publications. As discussed in the next section, this can be advantageous for gentle manipulation of cells.

### Fast, but gentle blood handling

Next, we investigated the performance of the acoustic mixer on human blood samples. We expected whole blood mixing to be more challenging because of cell-cell interactions in blood samples with increasing volume fraction of RBCs, or hematocrit (Hct). Normal Hct ranges from 36 to 45% for women and 40 to 50% for men, corresponding to viscosities of 5 to 100 cP at low shear rates ^28^. Hematocrit at birth is significantly greater, ranging from 42 to 65%. Here, we examined four different Hct (35, 45, 55, and 65%) thus covering the entire healthy human range. Human blood samples were obtained under local ethical approval from the Scottish National Blood Transfusion Service (contract #20∼11).

Upon arrival, samples of specific Hct values were made by adding or withdrawing plasma from these samples. 65 µL of each one of these artificially created samples was pipetted into the micromixer, followed by 65 µL PBS to assess the micromixer blood mixing capabilities. We applied the same image analysis technique as previously described (see Section 1) although images were acquired with digital microscope equipped with white light illumination (AM7013MZT, Dino-Lite) instead. Figure 3.a shows screenshots of the mixing experiment with the highest Hct blood sample (65%). Samples videos are available in SI Video 2. After 30 seconds, the human eye cannot distinguish any improvement in mixing. Image analysis (Figure 3. b) showed that after 30 seconds, the mixing performance reach almost 90% for all Hct concentrations. However, statistical analysis demonstrates that there is no significant difference between the 30 and 60 seconds mixing time point, as well as between some of the 10 and 60 seconds mixing. This may be due to the fact that in some experimental runs (4 out of 48 measurements), the mixing appeared to occur more quickly than usual. Some frequency drifts from the analogue function generator may have caused the intensity of the streaming to change and resulted in a faster mixing. After 60 seconds, mixing reaches its maximum (97.5% on average across all Hct concentrations). Importantly, from 30 seconds onwards, all Hct samples performed identically in terms of mixing (p-values non-significant in all cases). This demonstrates that the performance of the micromixer is unaffected by the increase in RBC volume fraction from 30 seconds onward. While the device does not reach the performance reported by Pourabed et al, it should be noted that Pourabed utilizes continuous flow driven by pumps, in addition to active acoustic mixing, and that here much larger volumes of blood are handled in a static manner. The advantage of a static volume is that no additional control of fluid, nor pump actuation is needed.

**Figure 3:**
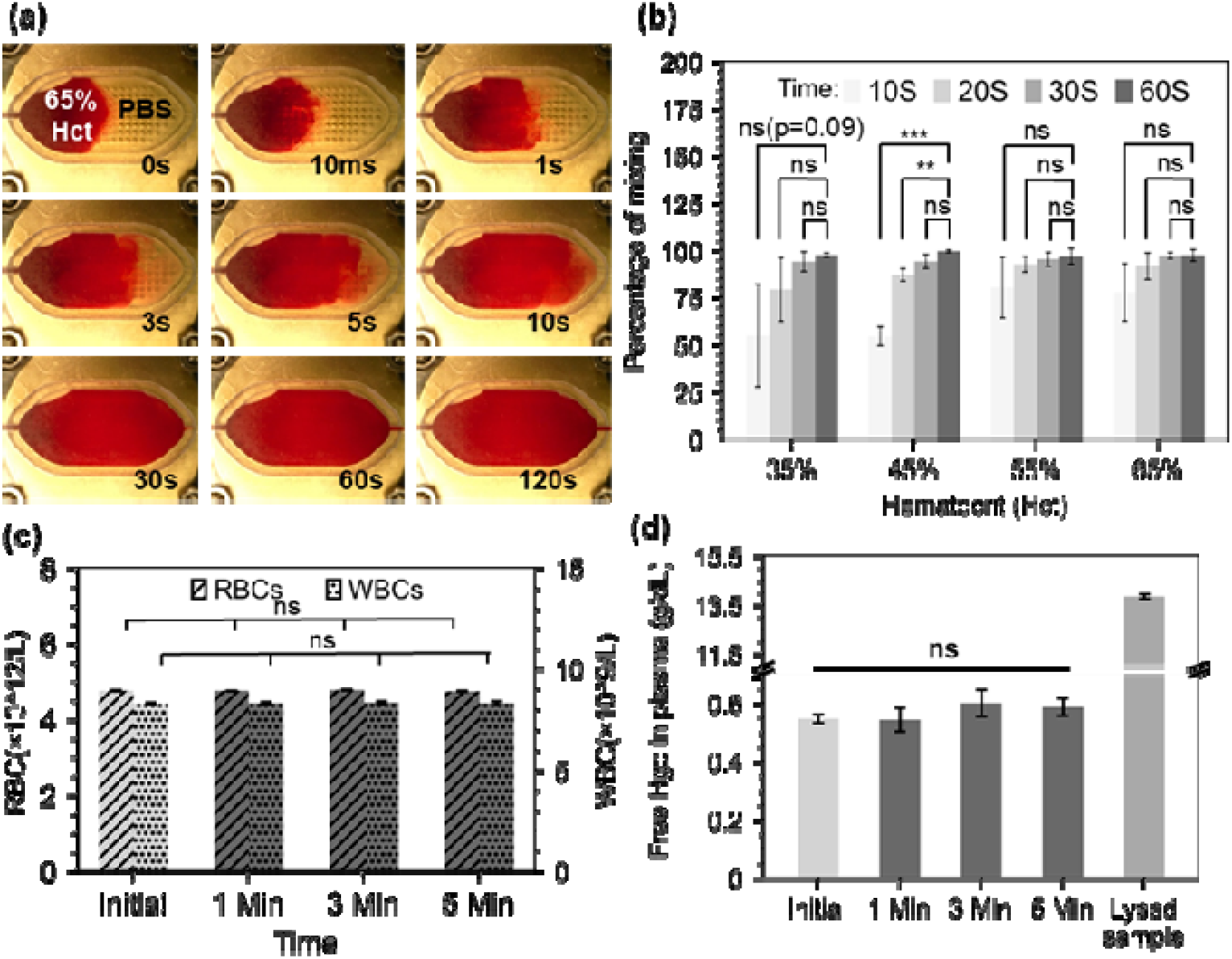
(a) Video screenshots of 65% Hct blood sample mixing with PBS at different time points (b) Percentage of mixing for various four Hct level at four time points. Statistical analysis: Student’s t-test between each time point and the 60 s time point. ns=non-significant. (c) RBCs and WBCs count for initial samples, after mixing of 45% Hct blood samples and lysed sample control. Statistical analysis: Annova (d) Free-Hgb for initial samples, after mixing of 45% Hct blood samples and lysed sample control. This set of results demonstrate the efficient agitation of blood sample, but absence of lysis for up to 5 min of continuous mixing. Statistical analysis: Annova. All data acquired in triplicate.

In laboratory testing, hemolysis, or the rupturing of RBCs, is a serious concern that can result in inaccurately raised potassium levels as well as a reduction in pH, pCO2, and pO2 values 29. Therefore, it is essential to verify that the microstreaming-based mixing is not harming RBCs while in operation and is not resulting in inaccurate analytical data. To evaluate that, whole blood samples (45% Hct) were processed on the mixer after 1, 3, or 5 minutes of mixing to see if the microstreaming agitation would cause blood lysis. A full blood count was obtained before and after mixing using a haematology analyser (Sysmex XP-300, Sysmex Corporation, Japan). The comparison between the initial blood count (RBC and WBC) and the processed blood samples count, demonst ate that no RBC or WBC damage occurred during the mixing up to 5 minutes (Figure 3.c).

When RBCs are damaged, they release their haemoglobin (Hgb) content onto the sample, hence free Hgb absorbance measurements in plasma can also be used to estimate the overall RBC lysis in the sample. This measurement cannot be directly taken with the Sysmex analyser since it has an in-built lysis step for total Hgb concentration measurement and a resolution of 0.1 g/dL limit, below the expected values for free Hgb in the absence of hemolysis. Therefore, collected samples were spun for 45 seconds at 13,400 rpm (Eppendorf MiniSpin Plus Centrifuge, Eppendorf, Germany), and absorbance measurements were measured on a spectrophotometer (Jenway 7315) on plasma after a 1:15 dilution with phosphate-buffered saline (PBS). A corrected absorbance by the Cripps method^30^ was calculated and a free Hgb value was obtained from an extrapolated standard curve (See SI Section 2). The results show no significant trace of excess Hgb on the processed samples up to 5 minutes (Figure 3.b), confirming that, unlike acoustofluidic structures designed for lysis, our acoustofluidic micromixer is gentle on whole blood samples^31,32^.

### The acoustofluidic mixer enables a 60 seconds lysis protocol

Since the micromixer does not infer hemolysis per se, additional cell lysis assistance is required to infer RBC lysis in human blood sample. Solutions for cell lysis, include chemical, mechanical, electrical, thermal, laser-based, and other lysis methods^33^.^33^. We chose chemical lysis as it is a standard laboratory method, and can be easily integrated into a lab-on-chip without additional parts necessitating electrical power. To evaluate the mixer performance in an RBC chemical lysis protocol, we loaded a finger-prick volume (21.6 μL) of human blood and 108 μL of Roche RBC Lysis Buffer (Roche, Merck, Germany) into the mixer for the on-chip protocol and mixed the blood and lysis buffer for 0 25, 0.5, 1, and 3 minutes. Control A and B are negative controls where the lysis buffer was replaced by PBS on bench and on-chip, respectively. Controls C and D are two positive bench controls which followed the exact manufacturer protocol of the above-mentioned RBC lysis buffer, but in control D, the initial volume of blood was 21.6 µL to match the on-chip volumes. Figure 4.a shows a series of spun (45 seconds at 13,400 rpm) samples: the original whole blood sample, the negative chip control (Control A), the sample under 1 min mixing, and the bench sample with matching dilution and volumes after 10 min mixing (Control D).

**Figure 4:**
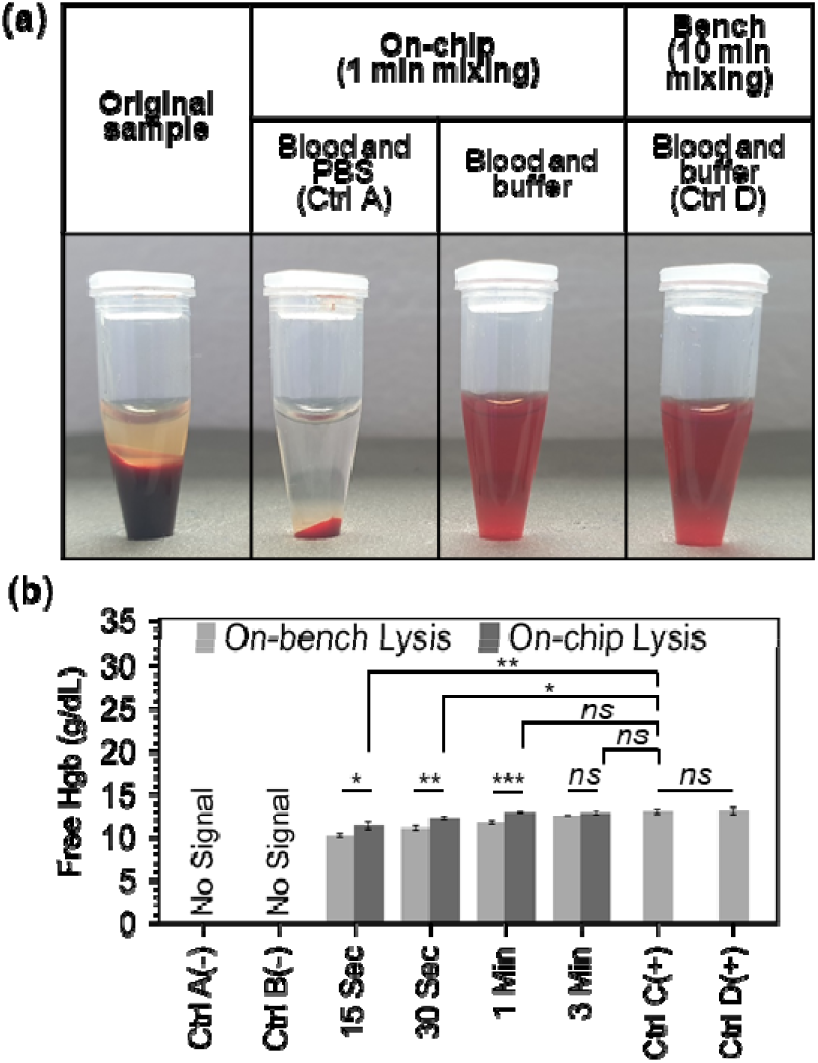
a) Photographs of centrifuged samples whole blood, blood mixed with PBS (1:5 v/v) and blood mixed with lysis buffer (1:5 v/v) following on-chip or bench mixing. b) Free Hgb measurement of on-chip and bench lysed samples. The negative controls, control A and control B, used 108 μL of PBS and 21.6 μL of blood on bench and chip, respectively. The positive controls C and D were both performed on bench w th 1 volume of blood diluted to 5 volume of buffer. Control C starting blood volume was 100 μL matching the manufacturer’s lowest volume requirement. Control D had a starting blood volume of 21.6 μL matching the chip volume. All controls use the manufacturer recommended time of 10 min. Statistical analysis: Student’s t-test. *, **, *** indicates p < 0.05, p ≤ 0.01, p ≤ 0.001. n.s. = non significance. All data acquired in triplicate.

Free Hgb absorbance measurements in plasma was used as previously described to estimate the overall RBC lysis in the sample. Our micromixer was able to lyse over 80% of RBCs in just 15 seconds, and 90% in 30 seconds. For complete RBC lysis, at least 1 minute of mixing was required (Figure 4.b). The micromixer outperformed the benchmark in terms of lysing RBCs at all time instances until complete lysis was accomplished in 1 min. This can be explained by the poor performance of hand inversion mixing (the manufacturer’s recommended mixing procedure) on small volumes. Additionally, to being superior to hand inversion, acoustofluidic mixing is also more reliable and is not as labour intensive.

Having demonstrated a rapid chemical-assisted lysis in just sixty seconds, we progressed to testing this on-chip protocol on healthy donor samples. Ten donor blood samples with hematocrit between 39.4% and 51.6% (respectively corresponding to 13.1 and 16.4 g/dL total Hgb) were acquired from the Scottish National Blood Transfusion Service (contract #20∼11). Each donor sample was processed following the lysis assay manufacturer protocol (100 μL Blood, 500 μL lysis buffer, 10 min) and following the on-chip lysis protocol as previously described for one minute. The total hemoglobin levels for the original samples were obtained from direct Sysmex reading (1:5 v/v dilution in PBS) while the hemoglobin levels for the processed samples were calculated as previously described (Figure 5.a). The free Hgb of the lysed samples was found to be highly consistent with the total Hgb level of the original samples, demonstrating complete lysis. In addition, the bench lysis and chip lysis exhibit a moderately high R-squared value of 0.885 in their correlation (Figure 5.b). Finally, all of the points are within the 95 percent margins of agreement for each lysis approach, according to a Bland-Altman analysis that we also used to assess the bias between the mean differences between the two lysis procedures (Figure 5.c). This demonstrates the excellent performance of our micromixer, which lyses RBCs in just one minute, or one tenth of the time recommended by the benchmark protocol.

**Figure 5:**
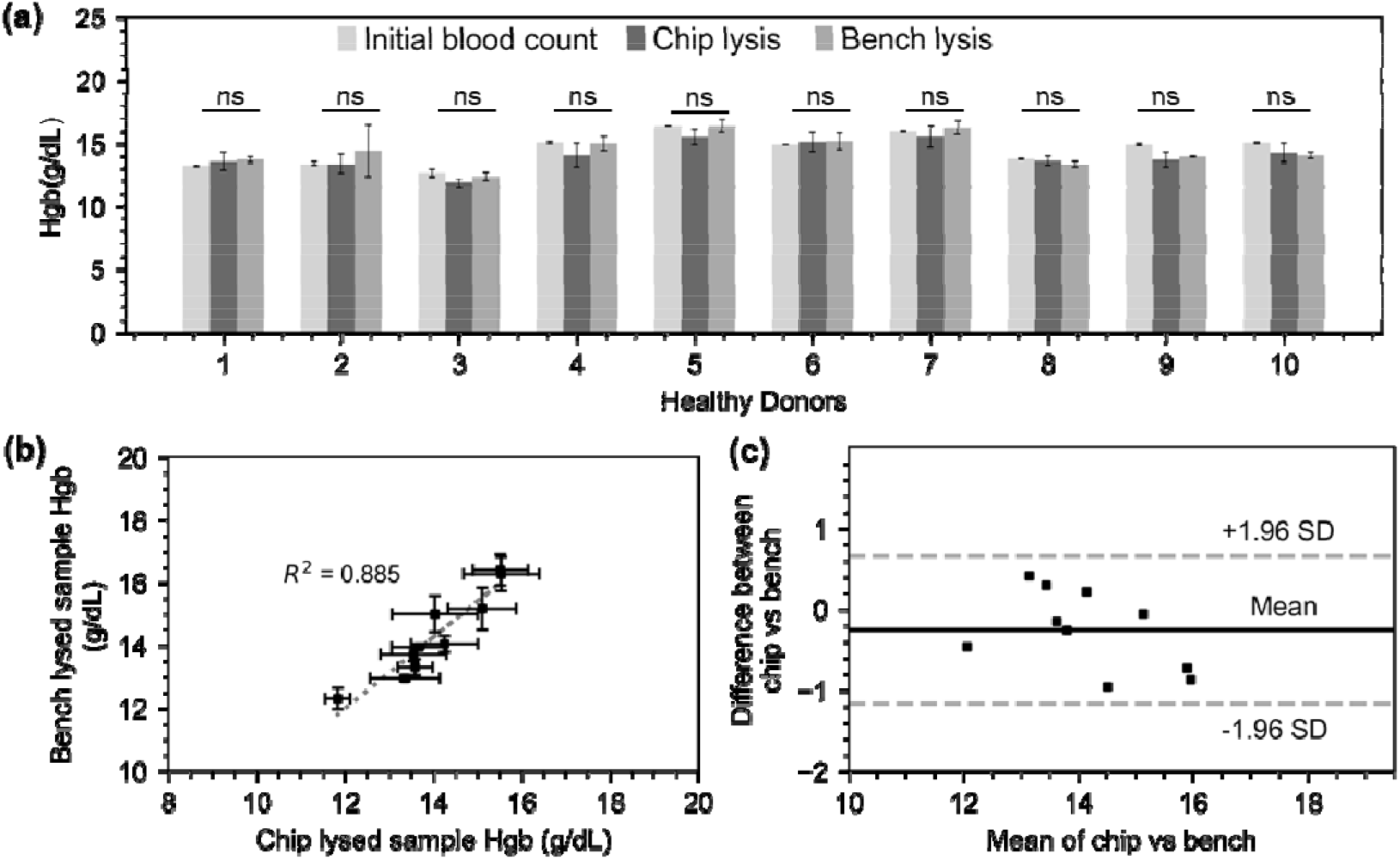
(a) Hgb levels on ten donor samples prior to processing, and after chip and bench processing. Statistics: Annova tests within samples showed no significant difference (b) correlation between bench and chip lysates. A linear regression model was fitted with an R^2^ value of 0.885. (c) Bland-Altman plot showing agreement between the hemoglobin values acquired on chip and bench lysates.

### An integrated finger-actuated sample preparation chip

Having demonstrated a rapid chemical-assisted lysis using the acoustofluidic mixer, we progressed to developing a finger-actuated device that would allow an untrained operator to obtain a blood lysate without any additional equipment apart from a blood collection device, and a generator for acoustic actuation.

Microfluidic-based procedures are sometimes made cumbersome and laborious by external connectors, tubing, and pumps. However, handling of finger-prick volumes of blood at the point-of-care should be easy and intuitive. A range of microfluidic devices allowing fluid pumping without the need for powered actuation or precise-pipetting, so-called ‘finger-actuated’ devices, were first introduced by Qiu et al in 2009^34^. This concept was further developed by a number of other groups, but most notably Juhwan Park and Je-Kyun Park, and demonstrated with the generation of gradients^35^, DNA extraction^36^, blood typing^37^ and blood-cross matching test^38^. The devices developed by Park and Park were manufactured in PDMS with standard SU8 based soft-lithography and displaced an average of 10 μL per stroke.

Our newly developed finger-actuated device uses a simpler, fast, manufacturing protocol based on laser cutting and solvent assisted bonding, which does not necessitate photolithography and cleanroom equipment. The device itself features seven layers allowing to connect the following features: **(i) a blood collection device interface**. The blood inlet geometry on the chip is designed specifically so that it can interface with the Minivette® POCT blood collection device, thus facilitating its use by avoiding unnecessary pipetting steps, but also providing enough actuation power to move the blood sample into a metering valve. **(ii) on-chip buffer storage**. We have embedded the lysis buffer on-chip in commercial blisters (150 µL Microfluidic Chip Shop, custom-filled with Roche RBC Lysis Buffer). Thanks to sharp needles engraved in layer 1, these blisters can be pierced upon a simple hand actuation which release the necessary buffer volume to lyse RBCs from whole blood samples (See SI Section 3 for details on fabrication and SI Video S3 for operation). **(iii) Metering and valving functions. (iii) Metering and valving functions**. Two turning valves enable the sequential addition of two pre-defined volumes of lysis buffer, the movement of the metered blood sample and the isolation of the mixing chamber during the lysis. Manual valves have the advantage of being completely power-free and of simple design, driving widespread adoption ^39–42^. A turning valve can also easily become a metering element, and we used one of these valves as such. The hollow section of the valve shaft contains 15 μL. Once blood is injected by actuating the blood collection device, the turning valve is rotated by 90 degrees and the second blister is pressed, flushing the content of the valve into the mixing chamber. The design and prototyping of these valves are described in SI Section 4, and a video of the valve operation in available is SI video 4. Once the lysis buffer and blood sample have been mixed and blood cells have been lysed, the lysate needs to be filtered out to remove RBC debris and other cells. **(iv) Embedded filter**. A Vivid™ asymmetric plasma separation membrane (Pall, GR grade, 12 mm &) sandwiched between layer three and five at layer four is used to filter the lysate. The entire content of the mixing chamber (130 µL) needs to be emptied and passed through the filters and ultimately delivered into a collection chamber. A video of the filter operation is available in SI Video 5 **(v) One-stroke pumping function**. In order to flush the blood lysate from the mixing chamber, a significant displacement of 130 μL is needed. The type of finger-actuated pumps previously proposed would not be suitable. By serendipity, a silicone cap commonly attached to disposable commercial vapes (ELF Bar 600) was found to have the perfect shape, flexibility and volume to displace the full content of the chamber in a single stroke (see SI Video 6). Using a blunt-tipped screw and a hammer, a 7.5 mm hole was cut into the top of the cap. This hole creates a favourable resistance ration and prevents backpressure from being created into the chamber and drawing back the fluid when the cap was released. For further precaution, the valve connected to the air cap can be closed after pressing the cap, to ensure full elimination of back pressure. The structure can be used as many times as required and does not need to be bonded onto the chip to function **(vi) Optical window**. In the final step of the on chip protocol, the filtered lysate is displaced into an optical window. This chamber is made by a cut-out through the full seven layers of the device and closed by two glass cover slips (American Scientific Products, M6045-1, 18×18 mm). The interfaced optical interrogation module can be used to examine the filtered hemolysate using spectrophotometry and other common optical instruments (not demonstrated in this work). The design and prototyping of each of the seven layers composing the blood lysate preparation chip are presented in Section SI 5.

To evaluate the repeatability of the fully integrated device, we fabricated a set of four identical devices, and processed aliquots of an additional human blood donor sample on this set of devices. A video of an operator demonstrating the complete on-chip protocol can be found in ESI Video 7. The filtered blood lysate was pipetted out of the chip after the full process, for characterisation. Figure 6.a shows the prototyped chip after the full process and before the lysate was pipetted out of the chip. For the bench control, the same donor blood sample was mixed with lysis buffer (1:5 v/v, n=4) following manufacturer instructions. The on chip and bench lysates were characterised by spectrophotometry as previously described. Since the dilution on the chip and on the bench are slightly different (valve metering volume 15 μL±2 μL and chamber volume around 130 μL), a relative absorbance measurement was performed. Figure 6.b shows the absorbance spectrum normalised to the mean (550-600 nm) for four on chip samples and four on bench samples. Except for one outlier, all of the on chip lysate absorbance spectrum perfectly match with those measured on the bench. Hgb typically exhibits high peaks at 576 nm; however, shifting of this peak, can occur either as a result of changes in chemical characteristics ^43^ or sample ageing ^44^. The on-chip outlier shows a similar spectra to the other samples, but is shifted with a peak at 574 nm. After carefully examination of the experimental videos, we suspect that a contaminant may have affected the sample properties. The comparison of the free Hgb absorbance values reveals a coefficient of variation (CV) of 15% for the on-chip lysate, compared to 4% for on-bench measurements Without the outlier, the CV of on-chip sample would be 3%, in-par with the bench control.

**Figure 6:**
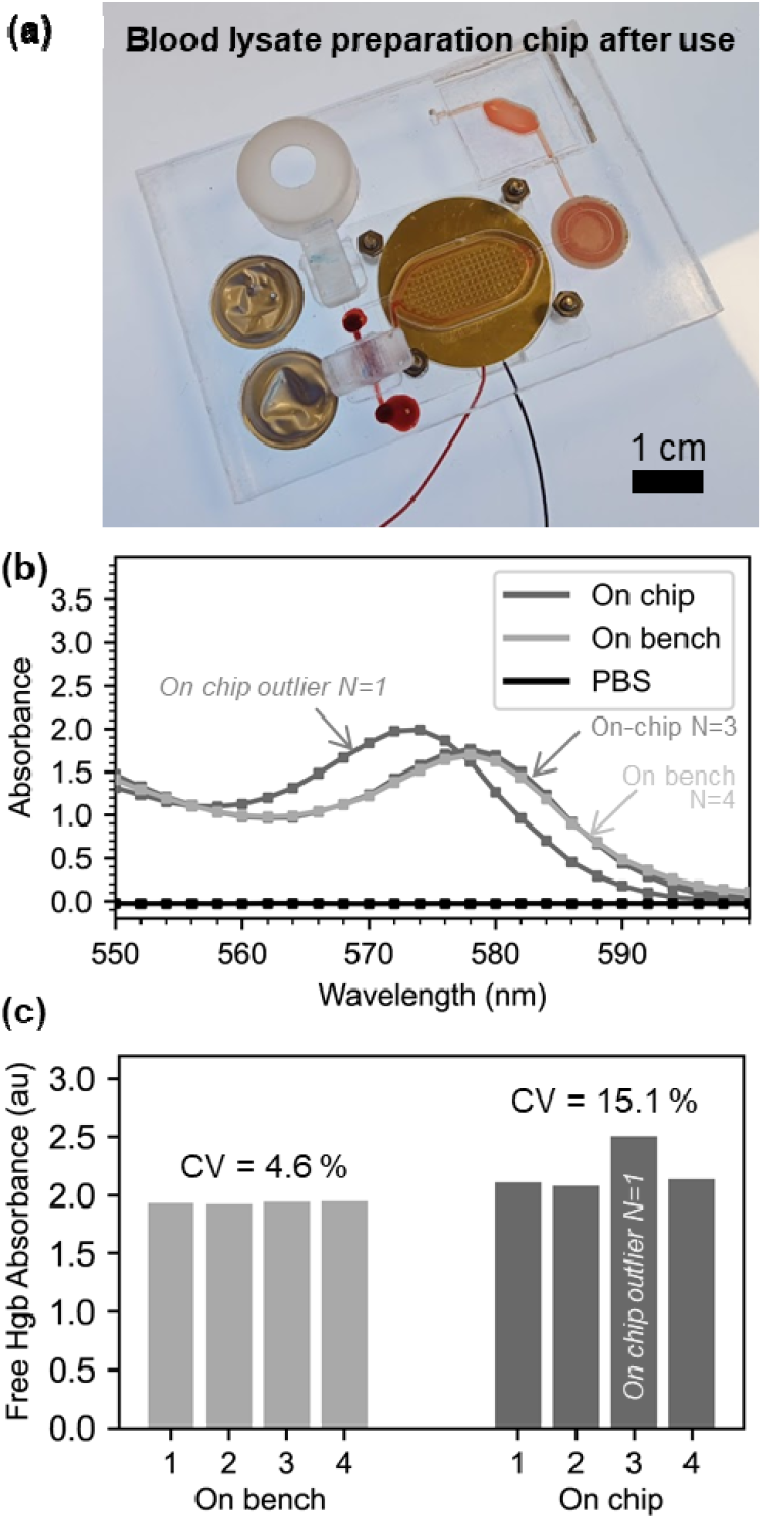
On-chip preparation of filtered blood lysate (a) Photograph of the integrated device after use. Components are arranges as illustrated in figure 1 (a) which identifies the operational parts of the chip. The scale bar is representing 1 cm. (b) Normalised 550-660 nm spectrum of on-chip and off-chip blood lysates. A mean normalisation was used. (c) Individual normalised free Hgb absorbance for each four on–chip and on bench sample (same donor)

These results demonstrate a range of useful features which enable effective sample preparation. The application of the findings and design features, are not limited to filtered blood lysate preparation. Similar features could be useful for the preparation of nucleic acids prior to polymerase chain reaction, for example. The device remains to be tested in conjunction with an optical detection system, or another kind of analytical system. This device is designed to be single-use and is prototyped in a fossil-derived plastic (PMMA). The single-use nature of the device brings convenience to the user, removes cross-contamination issues, and protects the user to harm with regards to potentially infectious samples, however this comes at a cost to the environment^45^. Improvements on both of these aspects will be the subject of future work. In particular, virgin PMMA could be replaced by recycled PMMA (Re-PMMA)^46^. We have demonstrated the compatibility of our solvent-assisted bonding protocol with Re-PMMA^47^. PMMA may also be substituted by sustainable materials such as PLA, or other sustainable biopolymers^48^. If the final application involved a high abundance biomarker, the assay may also be deployed in a more sustainable paper-based format.

## Conclusions

Handling of finger-prick volumes of blood should be easy and intuitive. Simple microfluidic-based procedures are sometimes made cumbersome and laborious by external connectors, tubing, and pumps. In this article, we have proposed a range of useful features for finger-actuated point-of-care integrated assays, all compatible with a straightforward, accessible and reliable prototyping technique. As far as we know some of these features have either never been reported (large volume, single-stroke (130 μL) finger pump) or never been previously integrated together (commercial blood collection device used as actuator and coupled with manual valve and blister reagent storage). In addition, an acoustofluidic mixer at the heart of the device provided gentle hemolysis-free acoustofluidic agitation of blood or efficient chemical-based complete lysis in less than 30s. The integration of these features was successfully demonstrated and benchmarked on ten human blood samples. Compared to the benchmarked manual RBC lysis assay, the use of the integrated chip was faster, and no pipetting was involved.

The features demonstrated here are using commercially available and cheap materials and protocols that do not necessitate expensive capital investment. The features presented can be used as a modular toolkit for point-of-care or point-of-use solutions. We hope they can help pave the way for more integrated and efficient on-chip protocols for the academic and industrial developments and facilitate exciting applications in human, veterinary or environmental health.

## Ethical statement

Human blood samples were obtained from the Scottish National Blood Transfusion Services under contract 20∼11. Local Ethical clearance was given by Heriot-Watt University EPS Ethics Committee.

## Author Contributions

M.E.H. and M.K.K conceived the study. A.J.C. conceived the concept of the hybrid acoustic micromixer. M.E.H. conceived the concept of the ‘one-stroke’ air cap. M.E.H. performed all experiments. W.N.M., S.R.K., R.M.C. and M. K. K. acquired funding. M.K.K. supervised the work and provided ethical clearance. M.E.H. and M.K.K. wrote the manuscript. All co-authors edited the manuscript.

## Acknowledgements

We acknowledge EPSRC 4MD programme EP/P027415/1 and EP/R00398X/1 for providing the funding necessary to complete this research. We are truly thankful to all our blood donors that kindly volunteered for this experiment.

