## Supplementary Information for "A microfluidic finger-actuated blood lysate preparation device enabled by rapid acoustofluidic mixing"

### Section 1. Method for the calculation of the mixing performance.

The video files (1280 x 960 pixels) were imported into Image J. An oval of width of 960 and height of 440 was placed on the first image. Next, the standard deviation of grey values (Analyse>set measurements>STD) was obtained on the first image was obtained in every stacks (Image>stacks>measure stacks). The file of standard deviations was saved and a manual note of the stack number at which the piezoelectric actuator was powered was made for every video. This was taken as the starting time point from then on.

The standard deviation file was opened in Excel and all data points before the starting stack previously defined were manually removed for all videos. A column with time was added (10 frames/sec video). The lowest standard deviation of the standard deviation column was identified, and the highest mixing standard deviation difference was calculated as lowest standard deviation subtracted by the initial standard deviation at starting time point. In each time point row, the initial standard deviation at the starting time point was subtracted from the recorded standard deviation. The percentage of mixing at each time point was calculated as the standard deviation at a particular time point divided by the highest standard deviation and multiplied by 100.

### Section 2. Method for the calculation of haemoglobin values

Sysmex hematology analyser was used to obtained complete blood counts, and total haemoglobin measurements on all original samples used in this study. However, the total haemoglobin measurement imposes an in-built lysis step, which prevents the measurements of free haemoglobin values which are indicative of hemolysis.

In order to derive free hemoglobin concentration values (g/dL) from absorbance measurement, a standard curve was created by correlating the total haemoglobin values obtained on the Sysmex hematology analyser and the free hgb absorbance calculated from the absorbance measurements from the spectrophotometer. Ten donor samples were received, and total hemoglobin was recorded on each samples with the Sysmex hematology analysers. The ten samples were then lysed according to the lysis assay protocol (Roche RBC Lysis Buffer), spun (45 seconds 12100×g) and 50 µL supernatant was diluted 1:15 in PBS prior to being measured on the Jenway spectrophotometer. Three more points have been added to the 10 donor samples in the standard curve, a plasma sample, an artificially made 10% Hct sample and another artificially made 72% Hct sample. The absorbance at 560 nm, 576 and 592 nm were obtained, and a free Hgb absorbance (Cripps, 1968, DOI: 10.1136/jcp.21.1.110) was calculated as:

$$Abs_{Free\ Hgb} = 2x\ (Abs_{576}) - (Abs_{576} + Abs_{592})$$

The free Hgb values following complete lysis were plotted against the total haemoglobin values from the Sysmex readings to obtain a standard curve (Figure S1).

The free hemoglobin concentration of unknown samples can be back-calculated from this linear regression fit, providing the same dilution factors are respected.

It can be noted that in the original paper by Cripps, plasma samples are spiked by known concentration of haemoglobin to derive a standard curve. Here the samples obtained from the SNBTS are a few days old, and exhibit some degree of hemolysis, therefore following the original Cripps method would not result in accurate correspondence between the original samples and the benchmark lysis method.

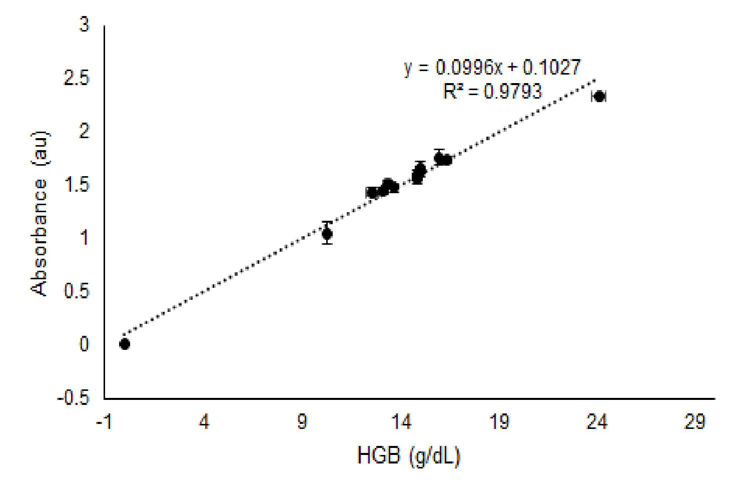


***Figure S1:*** *Standard curve for the calculation of free-Hgb from absorbance measurements.*

### Section 3 Integration of blister packs into the finger-actuated blood lysate preparation chip.

To integrate reagent-filled blisters, we faced the challenge of creating needle-like sharp structure using our laser cutting and solvent assisted bonding method (Liga et al 2016, https://doi.org/10.1007/s10404-016-1823-1). These blisters (here custom-filled by Microfluidic Chip Shop) are sold for integration into molded devices. For the design of the blister seat, and positioning of the needles we draw inspiration from Smith et al, 2016, DOI: 10.1007/s10404-016-1830-2.

The dimensions of the laser-engraved blister seat and needles are indicated in Figure S2A. We then engraved the blister seat around the needle on an Epilog Mini18 following the parameters in Figure S2B, creating the desired structures (Figure S2 C). Three photographs of an example blister assembling showing the blister assembled on a PMMA chip, and the blister crushed and liquid released (top and bottom view) demonstrating the piercing capability with a simple finger actuation.

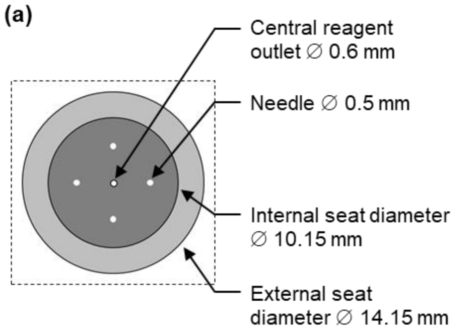


(b)

| CAD drawing | Laser Cut/Engrave parameters and depth | Photographs or resulting features |
|---|---|---|
| 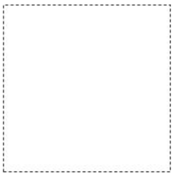 | *n.a.* | 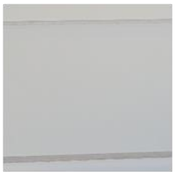 |
| 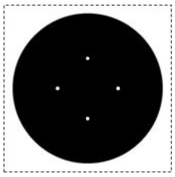 | Engrave<br>Speed: 22%<br>Power: 80%<br>Depth: 0.9±0.05 mm | 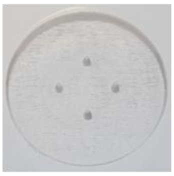 |
| 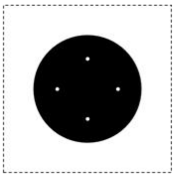 | Engrave<br>Speed: 22%<br>Power: 80%<br>Depth: 0.9±0.05 mm | 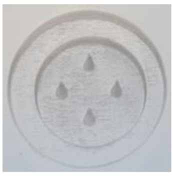 |
| 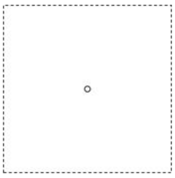 | Cut<br>Speed: 25%<br>Power: 90%<br>Depth: Cut through | 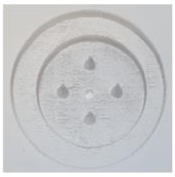 |

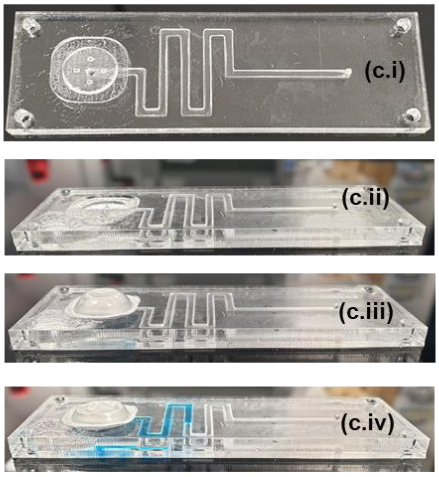


***Figure S2:*** *(a) Dimensions of the blister seat and piercing needles (b) Step-by-step guide to create blister seat and piercing needles (c) Series of photograph illustrating a test structure. (c.i) and (c.ii) Top and side view of structures prior to sealing of blister. (c.iii). Test structure with blister. (c.iv) Test structure after blister was pierced*

**Section 4. Creation of the manual turning valve.**

To enable the precise metering of the blood sample and we developed a reconfigurable turning valve module. The straightforward positioning, self-aligning features, and manual control of the turning valve makes the system user-friendly. This valve module takes advantage of the high-resolution manufacturing capabilities of stereolithography 3D printers and consists of two components: a valve shaft and a valve housing (SI Fig. 3a). The valve housing fits in a recess into the chip substrate.

The shaft was manufactured directly with a 3D printer using a clear resin (Form2, Formlabs). The housing was obtained by casting an elastomeric polymer (Sylgard 184, Dow Corning, standard mixing ratios) in a 3D printed mould using the same clear resin (SI Fig. 3b). By using these materials, the price of the valve (shaft and housing) is less than 0.30 USD.

The valve module can be integrated in microfluidic chips fabricated with the most common polymer manufacturing technologies enabling great versatility. This characteristics set a clear advantage over previous similar works, where only single channel control is possible (Guler et al, 2017; Bitsch et al 2006) and are only compatible with single microfluidic manufacturing technologies (Markov et al, 2010; Chan et al, 2016). The valve module can be plugged easily and directly into chips obtained from additive (stereolithography and fused deposition modelling), subtractive (laser cutting), and moulding (elastomer casting) microfluidic manufacturing technologies (SI Fig. 3c).

We demonstrated the performance of the turning valve by performing pressure burst tests in chips fabricated with each one of these technologies (Fig. 3d). Results indicate that the presented system can withstand pressures up to 1 MPa without noticeable leaking. In this present application, the pressures are minimal, and the manual valve shaft is therefore amply adequate.

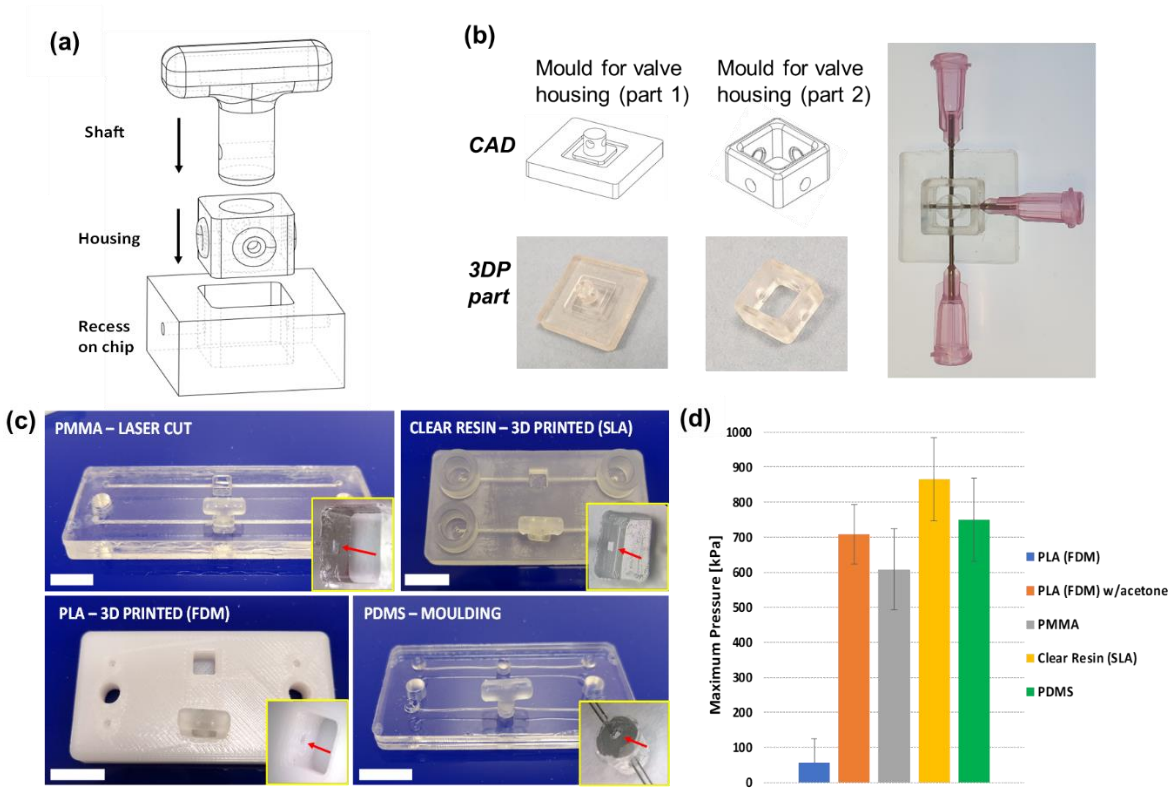


***Figure S3: (a)** 3D CAD representation of the valve structure, including the rigid 3D-printed shaft and soft silicone housing. **(b)** CAD and photographs of the mould for the silicone valve housing. b.i The use of a two part mould facilitate the demoulding. Two-part mould and needles in place. b.iii PDMS is poured into the mould. b.iv Demouldid valve housing. b.v assembled valve shaft and valve housing **(c)** Photographs illustrating the use of this valve on various protoytyping technologies: laser-cut PMMA, 3D printing (stereolithography), 3D (Fused Deposition modeling), PDMS moulding. **(d)** Maximum pressure withstood by the valve in closed mode by prototyping technology (N=3)*

### Section S5. Complete assembly of integrated blood lysate preparation chip.

Figure S4a presents each of the layer of the integrated blood lysate preparation chip. These layers were assembled by solvent assisted bonding technique described in Liga et al 2016 and in the video : https://www.youtube.com/watch?v=JzdkNnLQzMU After laser cutting on an Epilog Mini 18 (power and speed parameters in Fig S4b) layers were stacked together in the order presented Figure , and the pre-cut filter (Vivid GR) placed between layer 3 and 5 and ~160 µL of ethanol was pipetted on each of the layers. The stack was placed in a custom plate and into a Carver press at 3.5 metric tons, heated at 70 deg, for three minutes. After the devices had cooled down, the blisters were attached with the adhesive supplied by the manufacturer (Microfluidic Chip Shop), the valve housing, then valve shafts were inserted in the valve seats, the piezoelectric actuators were attached to the chip, and the glass cover slips (American Scientific Products, M6045-1, 18×18 mm), were attached with a pre-cut double-sided adhesive layer.

**(a)**

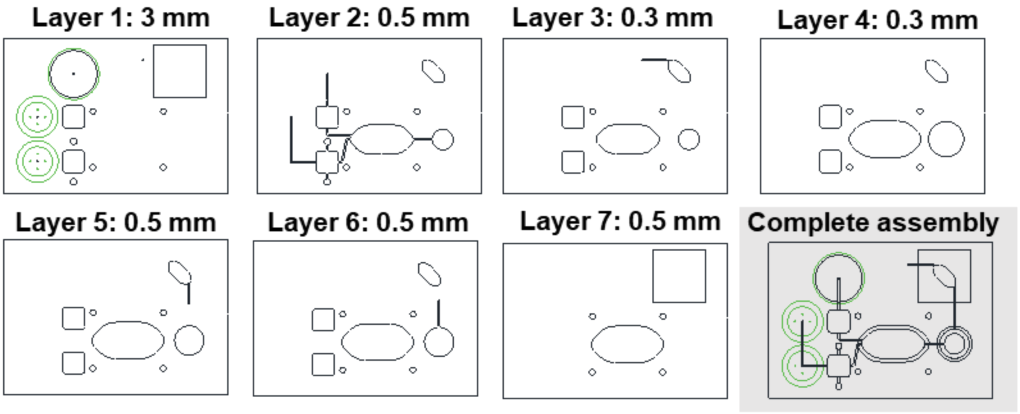


**(b)**

| | | Layer # | | | | | | |
|---|---|---|---|---|---|---|---|---|
| | | **1** | **2** | **3** | **4** | **5** | **6** | **7** |
| **Parameters** | **Vector cutting speed (%)** | 25 | 25 | 25 | 25 | 25 | 25 | 25 |
| | **Vector cutting power (%)** | 90 | 30 | 23 | 23 | 30 | 30 | 30 |
| | **Number of cutting passes** | 3 | 1 | 1 | 1 | 1 | 1 | 1 |
| | **Engraving speed (%)** | 22 | *n/a* | *n/a* | *n/a* | *n/a* | *n/a* | *n/a* |
| | **Engraving power (%)** | 80 | *n/a* | *n/a* | *n/a* | *n/a* | *n/a* | *n/a* |

***Figure S4: (a)*** *2D CAD of the seven layers of the integrated device, and overlay of the assembled layers. The total length of the chip is 76.2 mm (standard microscope slide length)* ***(b)*** *Vector cutting and engraving parameters for each layer*
